# Disease transmission mode has little effect on simulated canine rabies elimination

**DOI:** 10.1101/354175

**Authors:** Jessica K. R. Ward, Graham C. Smith, Giovanna Massei

## Abstract

Disease transmission in animal populations can be affected by density or by frequency of contacts between individuals, although many models assume linear density-dependent transmission. We used rabies and free-roaming dogs (*Canis familiaris*) in two Nepalese cities as a model system to explore the impact of linear and non-linear density-dependent disease transmission on disease elimination achieved through rabies vaccination or vaccination and sterilisation. Strongly non-linear transmission approximated realistic frequency dependent transmission. Free roaming dogs, abundant in many parts of the developing world, are responsible for most cases of rabies transmission to humans. Most models of rabies transmission assume that the disease transmission rate is linearly density-dependent, although a recent empirical study did not find evidence to support this assumption. Rabies vaccination, culling and dog sterilisation are employed to eliminate rabies or to reduce the numbers of human cases. We created a continuous-time deterministic compartmental model to describe rabies epidemiology within a dog population and to analyse the two modes of transmission. Under each transmission mode, we investigated the efficacy of dog vaccination and fertility control to eliminate rabies. To investigate the effect of dog density on disease control efforts, models were run for cities with high and low dog population densities using data from the Nepalese cities. The results showed that, at low population density, the amount of control effort required for disease elimination did not differ substantially between different transmission modes. At high population density, the effort required to achieve disease elimination was only higher for linear density-dependent than for non-linear transmission due to the exact value of the transmission rate, beta. The model suggests that although disease transmission mode may alter the impact of control on rabies elimination, this impact is relatively small and probably not relevant to disease management.

## 1 Introduction

Globally, canine rabies causes approximately 59,000 human deaths each year (Hampson et al., 2015), yet elimination of the disease is often still perceived as a low priority relative to other health programmes (Kaare et al., 2009). Over 3.7 million disability adjusted life years (DALYS) and economic losses of 8.6 billion USD per year are attributed to rabies (Hampson et al., 2015), although this may be a gross underestimation of the true burden of the disease (Belsare and Gompper, 2013).

Many countries in the developed world have greatly reduced or eliminated human rabies through vaccination and public education campaigns (Davlin and VonVille, 2012). Over 99% of rabies deaths occur in developing countries, where the disease is often endemic in domestic dog (*Canis familiaris*) populations (Hampson et al., 2009) and little control exists (Kaare et al., 2009). As more than 95% of the human deaths attributed to rabies occur as a result of bites by rabid dogs (Lembo and on behalf of the Partners for Rabies, 2012), eliminating the disease in dog populations is considered a high priority to reduce the global burden of canine-mediated human rabies (Meslin and Briggs, 2013).

In countries where large stray dog populations exist, the elimination of rabies has been the major driving force for dog population control and culling or sterilisation have been carried out to decrease the local abundance of dogs (Dalla Villa et al., 2010). Culling of free roaming dogs, that are often owned, is increasingly opposed by the public on the grounds of humaneness and effectiveness, and because of the environmental impact of toxicants (Hiby, 2013; Lembo et al., 2013). Culling may simply result in an increase in the demand for dogs within that community (Cleaveland et al., 2006). The World Health Organisation (WHO) do not recommend culling to control rabies as this can be ineffective and counter-productive, and instead promote mass vaccination campaigns as they have demonstrated marked success (WHO, 2013).

Empirical studies showed that canine rabies can be eliminated if 60-80% of the dog population is vaccinated (Cleaveland et al., 2003; Totton et al., 2010). This is in line with theoretical models that estimate that 70% vaccination coverage of a dog population is sufficient to eliminate rabies (Coleman and Dye, 1996; Totton et al., 2010) and this is the vaccination coverage recommended by the WHO (WHO, 2013), although this can be hard to obtain (Conan et al., 2015; Minyoo et al., 2015). This target is based on models that usually assume linear density-dependent disease transmission. However, a debate exists on whether the frequency of contacts between animals is more important than their density to drive disease transmission (McCallum et al., 2001; Ryder et al., 2007; Potapov et al., 2012; Morters et al., 2013). This would result in highly non-linear transmission and may therefore affect the target coverage at either high or low host density. Determining the mode of pathogen transmission is important for predicting the effect of disease control (McCallum et al., 2001) and rabies modelling is commonly used to inform control strategies (Sterner and Smith, 2006).

Many disease models assume that disease transmission is linearly related to host density; however, empirical support for this assumption is lacking (McCallum et al., 2001). There is growing evidence that simple classic models based on linear relationships between population density and contact rate are inappropriate (Barlow, 2000). Morters *et al.* (2013) failed to find a relationship between the reproductive potential of rabies (the average number of secondary infections produced by an infected individual) and dog density, and speculated that disease transmission might be frequency-dependent. True frequency-dependent transmission is unlikely, given that successful disease transmission depends on a number of different behaviours (e.g. social, territorial, sexual) and these are also adjusted by the rabies virus itself to increase the probability of onward transmission. Thus, here we assume a highly non-linear transmission as proposed by Barlow (Barlow, 1996) as a surrogate for ‘frequency-dependent’ transmission.

In linear density-dependent transmission the numbers of encounters between a susceptible and infected host per unit time will be linearly proportional to the density of infected hosts, as assumed by many previous rabies models (Anderson et al., 1981; Källén et al., 1985; Coyne et al., 1989; Smith and Wilkinson, 2003; Carroll et al., 2010). Under non-linear transmission, the probability that the susceptible host acquires an infection depends more on the number of contacts with other dogs which does not vary linearly with population density (McCallum et al., 2001; Sterner and Smith, 2006). Linear density and frequency-dependent transmission models are best thought of as extremes of transmission processes (Lloyd-Smith et al., 2005).

We used empirical data from two Nepalese cities with very different densities of dogs in a model to explore the effects of transmission mode on the time taken to eliminate rabies through vaccaintion. In Nepal approximately 200 people die of rabies each year (Acharya and Dhakal, 2015). As the main religions are Buddhism and Hinduism, which oppose the killing of animals (Kappeler and Wandeler, 1991; Kato et al., 2003), dog vaccination and fertility control have been used for the control of dog populations and rabies (Kappeler and Wandeler, 1991; Acharya and Dhakal, 2015). Where such control measures are used, the efforts required to implement these methods must be understood to improve control programmes, particularly because inadequate vaccination coverage may reinforce natural disease cycles, potentially increasing the disease problem (Hampson et al., 2007).

The aims of this study were 1) to determine what impacts the different transmission modes have on the control efforts required for rabies elimination in cities with different densities of dogs and 2) to explore how sterilisation would complement rabies vaccination to achieve disease elimination.

## 2 Methods

### Model description

A continuous time deterministic compartmental model was created to investigate the effects of disease transmission on the control of rabies in dogs. The model was adapted from Barlow (1996), Smith and Cheeseman (2002) and Hampson *et al.* (2007) and investigated the use of different disease transmission functions. The population was divided into three classes, Susceptible (*S*), dogs that had not been exposed to rabies, Exposed (*E*), which have been exposed to and are incubating disease, and Infectious (*I*), which are excreting virus and likely to be exhibiting clinical symptoms. The population dynamics are governed by a standard set of differential equations:

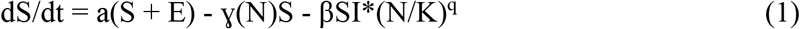

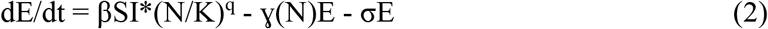

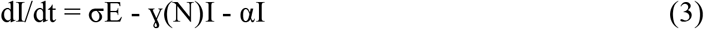

Where a = birth rate, ɣ = natural death rate, β= disease transmission coefficient, *N* = total population density, *K* = population carrying capacity, 1/σ = the mean incubation period, α = disease induced death rate and q = the degree of non-linear transmission (see below). The overall population is described by:

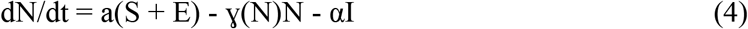

As the model uses continuous time, all rates are *per capita*, and *S, E* and *I* represent densities of each class, so *N* is the sum of *S, E* and *I*. Susceptible individuals enter the population through births at rate a(*S* + *E*). All classes suffer natural mortality as a function of density ɣ(*N*). Population growth therefore occurs at the rate *r* = a - ɣ(*N*). Mortality is assumed to be density-dependent where ɣ = *r/K* and thus increases as the population increases towards *K*, at which point it is equal to the birth rate.

Disease transmission is commonly represented by the term β*SI*. Here, we used the term β*SI* * (*N/K*)^*q*^ to investigate different modes of transmission. In this model *q* represents the degree of non-linearity in the contact rate: where *q* = 1, disease transmission occurs with linear density dependence (β*SI*). Where *q* <<1 a strongly convex-up relationship between contact rate and density occurs, representing realistic frequency-dependent disease transmission. At equilibrium there is no effect of *q*, however, both rabies and rabies management disturb the population from its equilibrium state and provide differing transmission rates.

Exposed animals become infectious at a rate σ*E* where 1/σ is the incubation period for the disease. Infectious animals suffer disease-induced mortality at a rate α*I*, where 1/α is the infectious period. The model assumed that no natural immunity to the disease exists within the population and that infected animals do not recover from rabies.

A vaccinated class (*V*) was added to the model as a form of rabies control. Susceptible individuals were vaccinated at a rate v*S*. It was assumed that exposed and infectious individuals do not respond to vaccination. Where vaccination does not last for the animals’ entire lifetime, immunity is lost at rate δ*V*, where 1/δ is the mean period of protection. The density of vaccinated animals is described by:

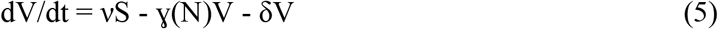

Where *N* is redefined to be *S + E + I + V*, and the whole population is described by:

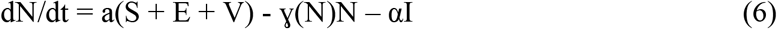

Since fertility control is known to improve rabies vaccination (Carroll et al., 2010), sterilisation was added to the model as a form of fertility control, with classes *S*_*f*_ *E*_*f*_ *I*_*f*_ and *V*_*f*_ representing infertile animals. Sterilisation of susceptible, exposed and vaccinated animals occurred at rate *p;* it was assumed that no infectious animals are collected for sterilisation. No animals in the sterilized classes could contribute to births as surgical sterilisation is effective for the animal’s entire lifetime. The sterile population can be described by the following equations with _*t*_ denoting the total (both the intact and sterile) population and *i* denoting only the intact population:

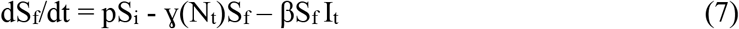

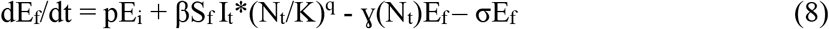

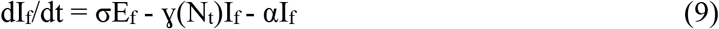

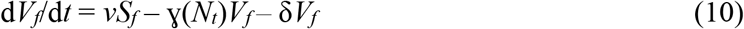

Therefore the overall sterile population can be described as:

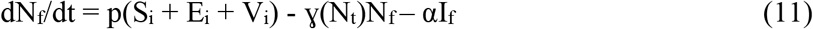

and the total population by:

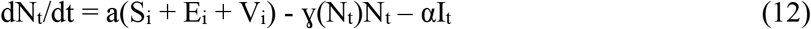

### Parameterisation

Data collected in Nepal were used to parameterise the model; where data were unavailable values were derived from the literature (Table 1). Birth rates were derived from data collected from household surveys (Massei et al., 2017) and from data provided by the local NonGovernment Organisation (NGO), the Himalayan Animal Rescue Trust (HART), based on records of surgical sterilisation carried out in twelve cities in Nepal. Household surveys showed that 74% of female dogs reproduced per year, with a mean litter size of 4.55 pups (SD 1.52, range 1-7 pups/litter). Thus birth rate is the proportion of female dogs that reproduce in any one month combined with average litter size. The male: female sex ratio of the studied population in the city of Pokhara was 1:0.52 (HART data) similar to previous reported values (Butler and Bingham, 2000; Kitala et al., 2001; Pal, 2001) giving an estimated monthly *per capita* birth rate of 0.098 for all populations.

As data on mortality were not available, mortality was assumed to be density-dependent. All density-dependence in the model was assumed to act through density-dependent mortality. Although density-dependence has been modelled acting on other processes (Barlow, 1996; Carroll et al., 2010) models with density-dependence acting only on mortality are often favoured as they produce a more conservative estimate (Kitala et al., 2002; Hampson et al., 2007; Carroll et al., 2010).

The degree of density dependence, *q*, chosen was 1 and 0.1 to represent biologically plausible extremes of density (linear) and frequency (non-linear) dependence.

**Table 1.** Parameter values used to inform the models at high and low population densities under frequency- and density- dependent disease transmission. β is the disease transmission rate, K is the carrying capacity, S, E I are the initial density of susceptible, infected and infectious dogs, 1/ σ the mean duration of incubation and 1/α the mean duration of infectiousness.

| Model location | Transmission mode | Parameter value |  |  |  |  |  |
| --- | --- | --- | --- | --- | --- | --- | --- |
| | | $\beta$ | $S$<br>$\text{km}^{-2}$ | $E$<br>$\text{km}^{-2}$ | $I$ $\text{km}^{-2}$ | $\sigma$<br>$\text{month}^{-1}$ | $\alpha$<br>$\text{month}^{-1}$ |
| **Kathmandu** | Linear | 0.05 | 518 | 0 | 1 | 1.04 | 5.35 |
| (K=525) | Non-linear | 0.03 | 518 | 0 | 1 | 1.04 | 5.35 |
| **Pokhara** | Linear | 0.55 | 30 | 0 | 1 | 1.04 | 5.35 |
| (K=35) | Non-linear | 0.45 | 30 | 0 | 1 | 1.04 | 5.35 |

Data from high-density and low-density populations of dogs in Nepal were used to represent extreme scenarios. The low-density population was modelled using data collected in Pokhara, where census data collected by HART gave a similar estimate to that of a previous report (Acharya and Dhakal, 2015), to produce an initial density of the 30 dogs km^−2^ for a stable population (estimated to be n=2,366) which we assumed was close to the carrying capacity, which we estimated at 35 dogs km^−2^. A high-density population was also modelled, based on published data of the dog population in Kathmandu, (Kakati, 2012) with an estimated density of 518 dogs km^−2^ (total population estimate 22,288) and again assumed to be close to carrying capacity which was estimated at 525 dogs km^−2^. There were no suitable data from these cities to parametrise the epidemiology.

The transmission parameter, *β*, needs to be estimated for each specific model (Caley et al., 2009). This needs to be set to simulate the same epidemiological outcome in both cities, regardless of the transmission function (since we ‘know’ the epidemiological trend over time but not the transmission function that drives it). Values therefore had to be adjusted until they produced damped oscillations of similar periodicity within each modelled population. Different initial values were necessary for high and low population density and for linear and non-linear transmission in order to produce realistic epidemics (Table 1) similar to those observed in dog populations (Hampson et al., 2007). Incubation and infectious period parameters were derived from literature values that reported natural rabies infections in dogs (Foggin, 1988; Fekadu, 1993; Coleman and Dye, 1996).

Disease control measures were implemented through rabies vaccination and fertility control (surgical sterilisation). Rates for both vaccination and fertility control were defined *per capita* (*v* in Eqn 5, *p* in Eqn’s 7 and 8) varying between 0 and 1 and thus approximating the proportion of the population treated in any one month. Numbers of treated individuals therefore varied with population density under the assumption that more resources would have been available in the higher density area. Due to the nature of the fertility control modelled (surgical sterilisation), it remained effective for life. Immunity from vaccinations is lost over time, most current vaccinations provide immunity for a minimum duration of 3 years (Schultz, 2006) and so the model included loss of immunity at a rate of δ=0.027 month^−1^.

The time taken for elimination of the disease within the population was investigated. A threshold for rabies elimination was set relative to the population size and elimination was considered to have been achieved when the number of infected individuals within the population was less than 1, and onward transmission therefore impossible. The equations assumed a closed population. Whilst the introduction of healthy animals would not affect the outcome, it is known that the movement of dogs incubating rabies do cause new outbreak in real life. However, once the population reaches the point of disease elimination, any introduction would not be successful as there would not be enough susceptible in the population until management stops and a new cohort of susceptibles are born.

For a combined approach (vaccination and fertility control) the level of control required to achieve rabies elimination within two years (24 months) was investigated. Models were run for both high and low population densities under both linear and non-linear transmission with nonlinear transmission causing a slow contact rate decline with density reduction (Figure 1).

All modelling was performed in STELLA^®^ version 9.1 (iSee Systems Inc., Lebanon, NH) and analyses were undertaken in R version 3.1.2 (R Core Team, 2014). Model verification was confirmed by a comparison with Carroll *et al.* (2010) and sensitivity of the model was determined by a one-at-a-time analysis.

**Figure 1.**
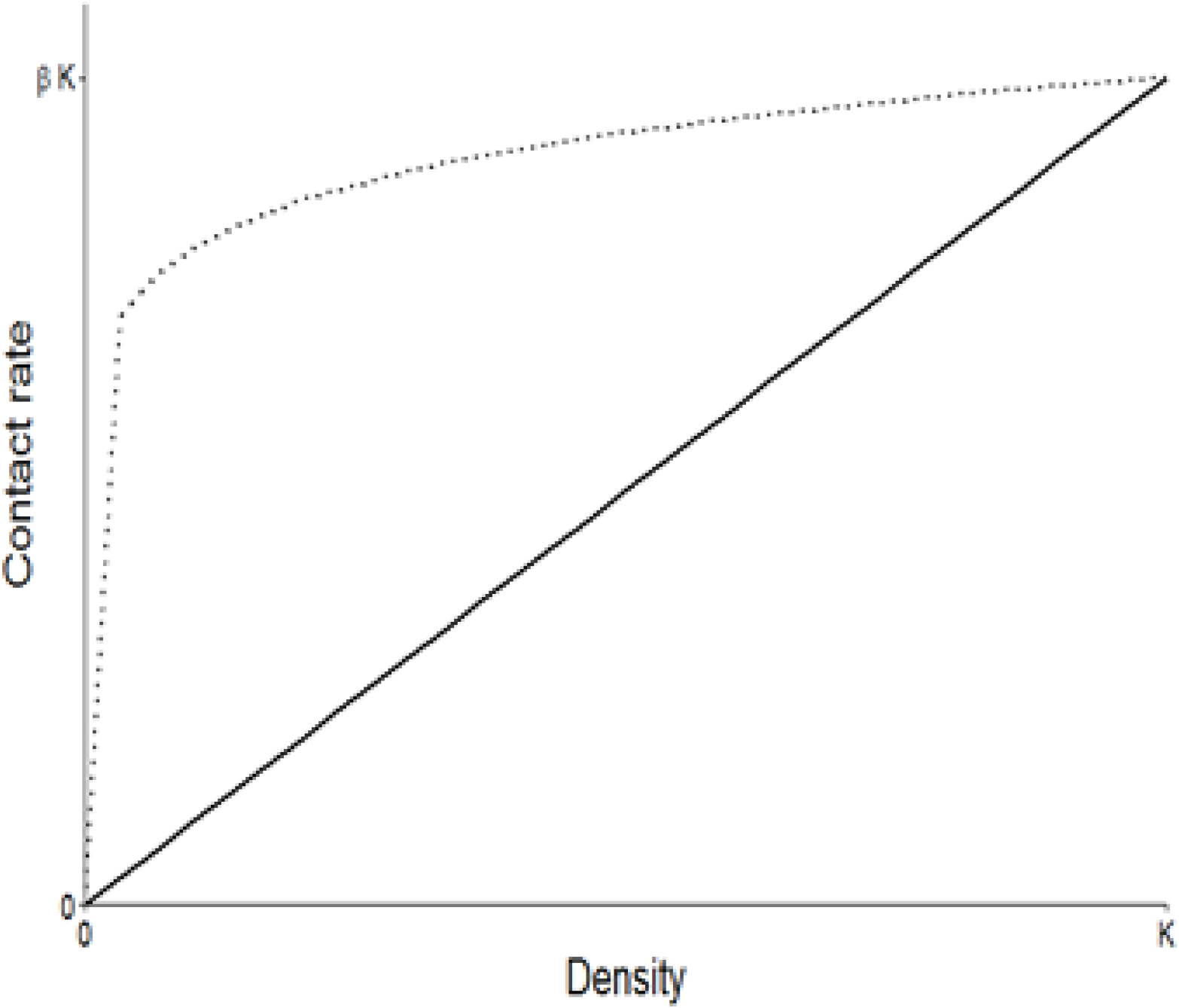
Adapted from Barlow (1996). Relationship between contact rate and density with contact rate increasing as the population density increases towards carrying capacity. The solid line indicates a linear relationship between contact rate and density and is represented by the transmission term β*SI*\*(*N/K*)^*q*^ where q=1. The dotted line indicates q=0.1 and represents an extremely non-linear ‘frequency’ dependent contact rate as used in the model. The transmission modes modelled represent extremes of system dynamics with the true transmission mode likely to be between the two extremes.

## 3 Results

By adjusting β, each version of the model produced a population which exhibited similar damped oscillations due to disease epidemics. Mean periodicity, measured as the time between peaks in population density, varied slightly between different disease transmission (27.3-33 months), and between high and low-density populations (27.3-30 months). In the models of non-linear transmission, the epidemics had a greater impact on population density, recovery of the population took longer and the average population density was reduced (Fig. 2a and b) when compared with the linear transmission model outputs.

Sensitivity analysis showed that parameters were more sensitive to change under the linear transmission model than the non-linear model. Birth rate was the most sensitive parameter with variations of +/−20% resulting in much greater changes in the average number of infected individuals over a 30 month period than any other parameter.

**Figure 2.**
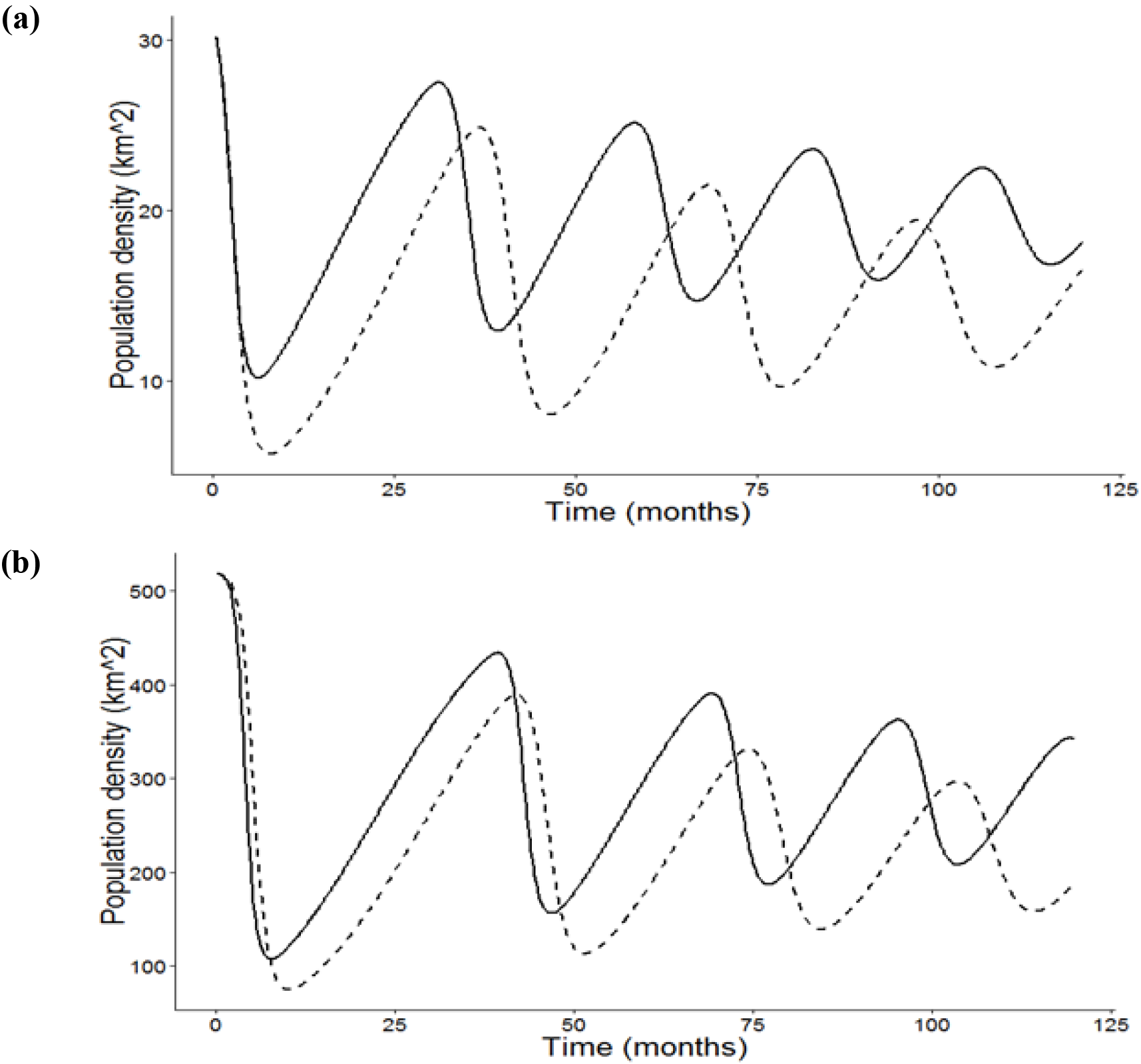
Model output showing the behaviour of rabies epidemics and their effect on domestic dog population densities in (**a**) low-density and (**b**) high-density populations based on data from Pokhara and Kathmandu. Solid lines indicate linear density-dependent disease transmission; dashed lines indicate strongly non-linear disease transmission. Under the latter transmission model epidemics are larger and the population takes longer to recover than under the linear density-dependent model.

### Disease elimination based on rabies vaccination and transmission mode

At a low dog population density, a minimum vaccination rate of 0.2 month^−1^ was required to achieve disease elimination regardless of transmission mode (Fig. 3a) and this was always achieved within two years. Under linear transmission, increasing the vaccination rate gave a minimum and maximum elimination time of 10.25 months and 16.5 months respectively. Nonlinear transmission gave minimum and maximum elimination times of 10.25 months and 22.75 months. Minimum elimination times were the same for both transmission models, however maximum elimination times varied between models with elimination being achieved faster under the linear transmission model. This preference was not sensitive to β.

**Figure 3.**
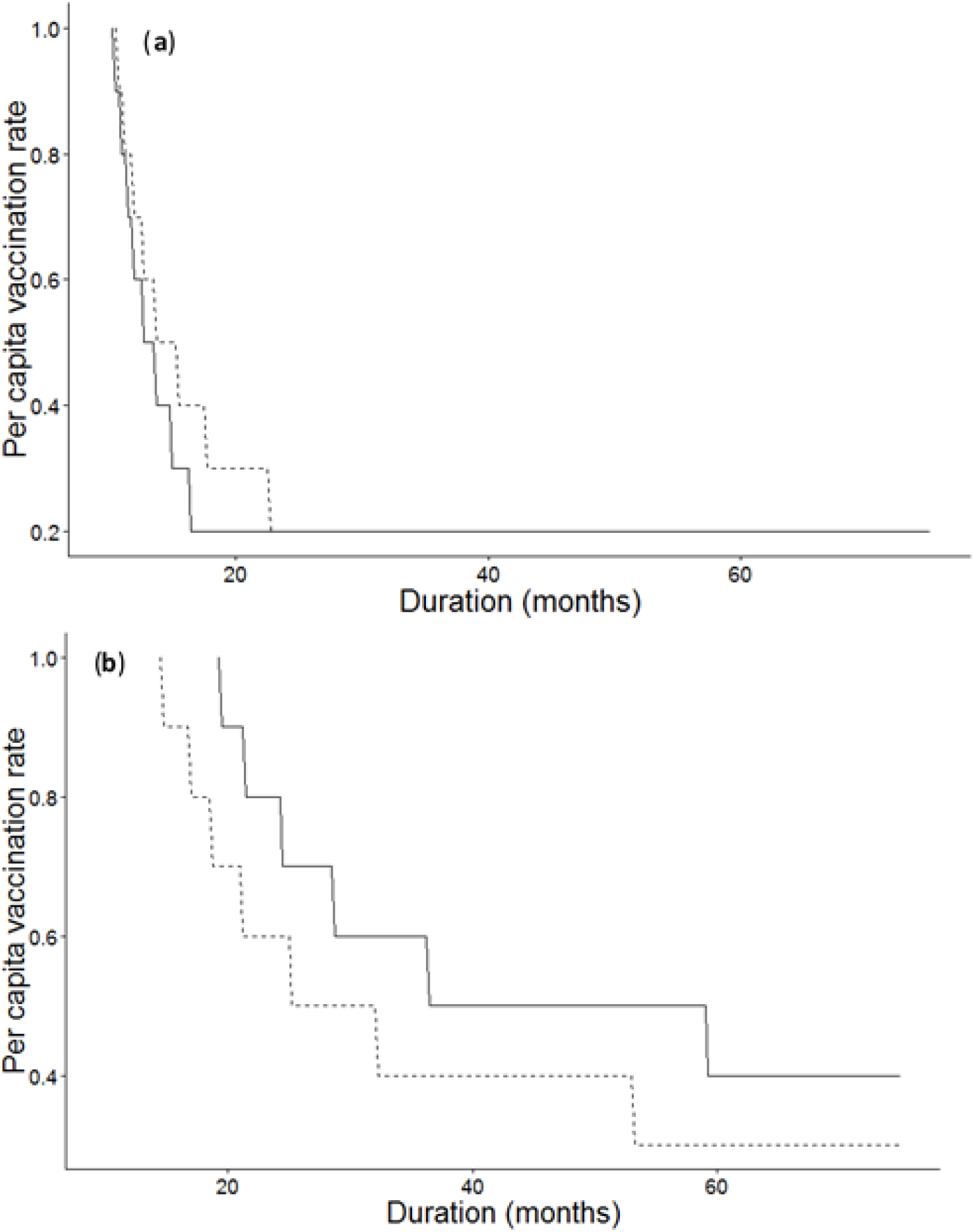
Time (months) taken to achieve disease elimination in (**a**) low-density and (**b**) high-density populations. Solid lines indicate linear density-dependent disease transmission; dashed lines indicate non-linear ‘frequency-dependent’ disease transmission. For each model the area under the curve indicates treatment levels at which rabies is not eliminated from the population, area above the curve indicates treatment at levels that are sufficiently high enough to achieve rabies elimination. The effect of transmission mode on rabies control is different under the different density models, in the low-density population differences between elimination times achieved under each transmission mode are minimal but differences seen are greater under the high-density model.

At high dog population density, a minimum vaccination rate of 0.3 month^−1^ was required under the non-linear model to achieve elimination (Fig. 3b) and under the linear model a minimum vaccination rate of 0.4 month^−1^ was required. At a higher density, this higher rate of vaccination means that a substantially higher absolute number of animals need to be treated to achieve disease elimination. Under linear transmission minimum and maximum disease elimination times were 18 and 59.25 months respectively. At high density, elimination was achieved faster under non-linear transmission than linear transmission, with elimination times ranging between 14.75 and 53.25 months in the non-linear model. However, the sensitivity analysis showed that small changes to *β* could alter this preference, indicating that the absolute value of β was more important than the mode of transmission.

**Figure 4.**
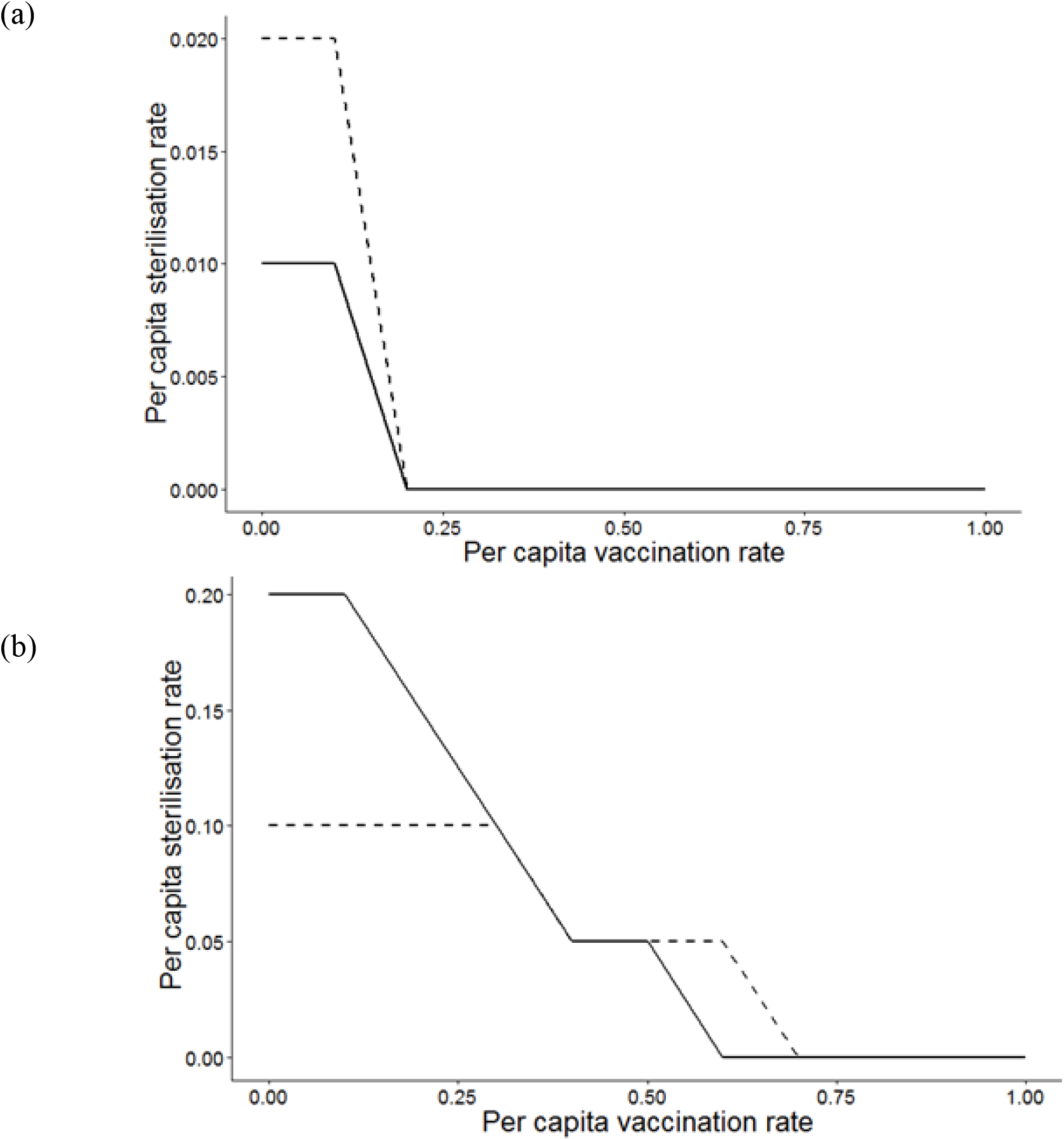
Combinations of sterilisation and vaccination rates required to achieve disease elimination within 24 months of continuous control in (**a**) low-density and (**b**) high-density populations. Solid lines represent linear density-dependent transmission in the model; dashed lines represent non-linear ‘frequency-dependent’ disease transmission in the model. For each mode the area under the curve represents treatment rates that are insufficient to eliminate disease, area above the curve indicates treatment (vaccination and sterilisation) levels that exceed the minimum values required to achieve disease elimination.

### Disease elimination based on rabies vaccination and fertility control and transmission mode

In the low-density population, disease was easily eliminated within two years of combined rabies vaccination and fertility control (Fig. 4a). Fertility control was necessary as an additional strategy at vaccination rates lower than 0.2 month^−1^. On its own, the minimum fertility control rates required to eliminate disease from the population within two years were 0.01 month^−1^ under linear transmission, and 0.02 month^−1^ under non-linear transmission.

In high density populations, the minimum vaccination rate required to achieve disease elimination within the 24 month time period in the absence of additional fertility control was 0.7 month^−1^ under linear transmission and 0.6 month^−1^ under non-linear transmission (Fig. 4b). At lower vaccination rates higher compensatory sterilisation rates were required to achieve disease elimination under linear transmission than non-linear transmission. Where fertility control was implemented in the absence of vaccination minimum control rates required to achieve elimination were an order of magnitude higher than in the low density population: 0.1 month^−1^ and 0.2 month^−1^ under non-linear and linear transmission respectively.

Since in the above simulations the optimum control switched from linear to non-linear transmission as density increased we ran further simulations to investigate this. In all cases the linear transmission model had higher values of β to obtain a similar epidemiological cycle. We therefore tested different densities and different values for β. At low density similar values of *β* still led to a preference for the linear model, rather than the non-linear model. As density increased, the necessary greater absolute difference in β between the models meant that the preference was always for the non-linear transmission, but small changes in β could change this preference.

## 4 Discussion

There is considerable debate on how disease control may be affected by disease transmission mode (Barlow, 1996; McCallum et al., 2001; Sterner and Smith, 2006). The model used was site specific and based on data collected in two Nepalese cities, Pokhara and Kathmandu. The large variation in recorded dog density between these cities provided an ideal scenario to test assumptions concerning the transmission processes.

The assumed pregnancy rate used in this model was higher than previous published estimates of between 47.5% and 54% females giving birth in any one year (Kitala et al., 2002; Reece et al., 2008). These differences may be explained by the fact that, for instance in the Indian city of Jaipur, bitches were not caught for sterilisation when heavily pregnant or lactating: thus the proportion of females found pregnant during surgical sterilisation might have not reflected the true proportion of dogs reproducing (Reece et al., 2008). The sex ratio estimated from the data 1:0.52 males: females is also lower than previous published estimates of 1:0.67-1:0.79 males: females (Butler and Bingham, 2000; Kitala et al., 2001; Pal, 2001). However, according to the sensitivity analysis, this had a minimal effect on the model. Male dogs in Nepal and neighbouring countries are known to be preferred for their use as guard dogs and females are killed in greater numbers than males as they produce unwanted litters or attract males when in oestrus (Beran and Frith, 1988; Butler and Bingham, 2000; Matter et al., 2000; Totton et al., 2010; Acharya and Dhakal, 2015) thus we are confident in the highly skewed estimate used in the model.

Mortality in the model was assumed to be density-dependent. Although density-dependence acting on mortality and natality can impact upon disease control (Barlow, 1996), the model was run for both high and low-density populations to minimise this effect. An inherent assumption of non-spatial linear density-dependent models is homogeneous mixing within the population (Anderson, 1982). However, non-random mixing may result in a contact frequency between hosts that is independent of density (Begon et al., 1999). Density independent contact could be affected by the potential alteration in behaviour associated with rabies (Sterner and Smith, 2006). Substantial changes in the social and or spatial behaviour of infected or infectious dogs might alter the results of these types of models.

The model assumed a closed population with no dogs being removed from or brought into the population and thus low levels of sterilisation could be effective. However, dogs are often traded into and out of areas according to human demand (Morters et al., 2014). The immigration of dogs, currently not accounted for in the model, needs to be taken into account when evaluating the use of vaccination, and fertility control, as this will decrease the control efficacy.

Although previous rabies models have largely assumed disease transmission was linearly density-dependent (Anderson and May, 1981; Smith and Cheeseman, 2002; Sterner and Smith, 2006), limited field studies, on other diseases, suggest that transmission may be more closely similar to frequency-dependent (Begon et al., 1998; Ramsey et al., 2002; Begon et al., 2003; McCallum et al., 2009). Rabies is spread by direct contact associated with behaviours such as conspecific aggression, mating and maternal protection of puppies: thus contact rate may vary through the year and with density (McCallum et al., 2009). With numerous behaviours influencing contact rates the true transmission mode of the disease is likely to depend on both frequency of contacts and density-of animals (Begon et al., 1998; Ryder et al., 2007). Morters *et al.* (2013) find some evidence for rabies in dogs that the relationship is at best a weak one with density. Models representing either linear density-dependent or fully frequency-dependent disease transmission can be thought of as representing two extremes. Disease control is therefore most likely if the predicted effort to achieve disease elimination under both linear and non-linear models is exceeded.

Assuming density-dependent transmission gives rise to a threshold density below which the disease cannot be maintained within the population (Anderson et al., 1981; Lloyd-Smith et al., 2005; Sterner and Smith, 2006; Morters et al., 2013) and such thresholds are thought to vary across ecosystems and habitats (Begon et al., 1999; Lloyd-Smith et al., 2005; Sterner and Smith, 2006). Should disease transmission be entirely frequency-dependent, then a threshold population density would not occur (Lloyd-Smith et al., 2005; Ryder et al., 2007), although a more realistic non-linear density dependent model would still retain a threshold, albeit much lower. Control strategies focussed on reducing the susceptible population to below such thresholds through lethal techniques could therefore be misguided. Where a disease has an element of frequency dependence in its transmission a single infected individual retains the potential to infect numerous individuals regardless of the overall population size.

This work indicates that disease control is relatively harder in high density populations, with a higher rate, or proportion of the population, needing to be treated to obtain disease control. The mode of disease transmission that resulted in easier disease management also appeared to switch as host density increased. However, sensitivity analysis has shown that the absolute value of β used is an important driver in this preference. The determination of a linear or non-linear mode of transmission is less important, as long as the model reproduces realistic temporal patterns.

Therefore host density appeared to be much more important than the mode of disease transmission. Additionally, simulating realistic temporal trends may be more important than the mode of transmission. As there was relatively little difference between the results of the two disease transmission modes, control programmes should aim to exceed the threshold for control from both linear and non-linear models.

At least for canine rabies, the transmission mode causes little difference in the amount of control required to achieve disease elimination, although a slight reduction in the effect of sterilisation was seen under non-linear transmission, in agreement with Barlow (Barlow, 1996). This study confirms that adding fertility control to vaccination will reduce the effort required to eliminate a disease by reducing the overall proportion of animals that must be treated, and this is in agreement with Carroll *et al.* (2010). Focusing disease control on combined vaccination and fertility control should be encouraged as the efficacy of such programmes is less affected by disease transmission mode.

## 5 Acknowledgements

Thanks to Dr Sarah Jayne Boulton and A. Higgins for their comments on the initial draft and to Barbara Webb, Jim Pearson and HART for providing data on Pokhara. This research did not receive any specific grant from funding agencies in the public, commercial, or not-for-profit sectors.

